# Attenuation of neural responses in subthalamic nucleus during internally guided voluntary movements in Parkinson’s disease

**DOI:** 10.1101/2022.09.09.507256

**Authors:** Veronika Filyushkina, Elena Belova, Svetlana Usova, Alexey Tomskiy, Alexey Sedov

## Abstract

The proposed models of segregated functional loops describe the organization of motor control over externally triggered (ET) and internally guided (IG) movements. The dopamine deficiency in Parkinson’s disease (PD) is considered to cause a disturbance in the functional loop regulating IG movements. At the same time, the neural mechanisms of movement performance and the role of basal ganglia in motor control remains unclear.

The aim of this study was to compare neuronal responses in subthalamic nucleus (STN) during ET and IG movements in PD. We found and analyzed 26 sensitive neurons in 12 PD patients who underwent surgery for implantation of electrodes for deep brain stimulation. We also analyzed the local field potentials (LFP) of the STN of 6 patients in the postoperative period. Patients were asked to perform voluntary movements (clenching and unclenching the fist) evoked by verbal command (ET) or self-initiated (IG).

We showed heterogeneity of neuronal responses and did not find sensitive neurons associated with only one type of movement. Most cells were characterized by leading responses, indicating that the STN has an important role in movement initiation. At the same time, we found attenuation of motor responses during IG movement versus stable responses during ET movemements. LFP analysis also showed attenuation of beta desynchronization during multiple IG movements.

We propose that stable neuronal response to ET movements is associated with reboot of the motor program for each movement, while attenuation of responses to IG movement is associated with single motor program launching for multiple movements.

## 1 Introduction

Parkinson’s disease (PD) is a neurodegenerative disorder caused by the death of dopaminergic neurons in the substantia nigra pars compacta. The lack of dopamine leads to malfunctioning of neural loops inside the basal ganglia (BG) network and, consequently, to excessive inhibition of the motor thalamus, projecting directly to the motor cortex (Alexander et al., 1986). The increased inhibition leads to hypokinetic signs in patients with PD (Jahanshahi et al., 1995; Buhmann et al., 2003; Nambu, 2008).

According to motor control models, self-initiated (internally guided, IG) movements are controlled predominantly by the basal ganglia-thalamus-motor cortex (BGTM) loop, while externally triggered (ET) movements are promoted mainly by a cerebellar-cortical (CC) loop (Cunnington et al., 2002; Cerasa et al., 2006; Taniwaki et al., 2006, 2013; Purzner et al., 2007; Hackney et al., 2015).

The BGTM loop encompasses motor areas of the cortex such as the supplementary motor area (SMA), ventral premotor cortex (PMv) and sensorimotor cortex (SMC) along with several subcortical structures - the putamen (Put), the globus pallidus (GP) and the ventrolateral nucleus of the thalamus (VL) (DeLong, 1990; Hasegawa et al., 2022). The cerebellar-cortical loop consists of the motor parts of the cortex (SMC or PMv) receiving projections from the anterior lobe of the cerebellum (CB) through the dentate nucleus of the cerebellum (DN) and the ventral posterior lateral (VPL) nucleus of the thalamus, while the cerebellum receives inputs from the motor cortex through the pontine nucleus (Kelly and Strick, 2003).

Dopamine deficiency in PD causes an impairment of the BGTM functional loop, which regulates predominantly internally guided movements, whereas emerging external signals may facilitate motor execution (Martin, 1967). The possible explanation for this is that these functional circuits are segregated and unaffected cerebellar-cortical loop may be mainly involved in the implementation of externally triggered movements opposed to internally guided ones (Taniwaki et al., 2006, 2013; Hackney et al., 2015). Another hypothesis based on functional MRI studies proposed that in normal condition ET and IG movements are primarily processed through СС и BGTM circuits, respectively, with the recruitment second pathway. In PD ET tasks are primarily encoded in the CC circuitry including the BGTM circuitry. However, IG movements in PD neither adequately activate the BGTM pathway, nor cause adequate recruitment of the cerebellar-cortical pathway. As a result, both pathways display decreased activity (Lewis et al., 2007).

Most of the studies dealing with ET and IG movements focus on the research of anatomically and functionally segregated brain pathways contributing to motor control for these types of movements (Jahanshahi et al., 1995; Cunnington et al., 2002; François-Brosseau et al., 2009; Filyushkina et al., 2019). Sparse electrophysiological studies on this topic showed that the BGTM is involved in the preparation of both self-paced and externally cued movements while the CTC pathway is involved in the preparation of self-paced but not externally cued movements (Purzner et al., 2007). Only externally-cued movements induced a pro-kinetic event-related beta-desynchronization, whereas beta-oscillations were continuously suppressed during self-paced movements (Bichsel et al., 2018).

The subthalamic nucleus (STN), being an important part of the basal ganglia network, projects to the external and internal pallidum, the modulatory and output nuclei of the BG, respectively, and plays an indispensable role in controlling voluntary movements (Purzner et al., 2007; Bichsel et al., 2018; Hasegawa et al., 2022).

However, the precise mechanisms for movement control within the STN remains unclear as well as the neural underpinnings for IG and ET movements and the pathophysiological mechanisms that disrupt motor function in PD. Here we aim to study the neuronal mechanisms of motor control in STN at the level of single unit activity and local field potentials in patients with PD performing the same ET and IG movements.

## 2 Method

### Data collection

The study of the activity of STN neurons responding to motor tests included 12 patients (6 females and 6 males, average age 53.1±8.4, mean disease duration was 9.9±1.9 years) (Table 1). Local field potentials were studied in the postoperative period in 6 patients (5 female, 1 male, average age 53.8±8.2, mean disease duration was 13.7±4.5 years), who underwent lead externalization (Table 1). The disease severity was estimated using the unified PD rating scale (UPDRS)-III, without levodopa administration (OFF-state). For patients in the intraoperative group, the UPDRS-III scores were assessed before operation and ranged from 18 to 78 points, for postoperative group, the UPDRS-III scores were assessed just before the start of recordings and ranged from 26 to 57 points. All patients were withdrawn from medications at least 12 hours before the start of the recordings.

**Table 1.**
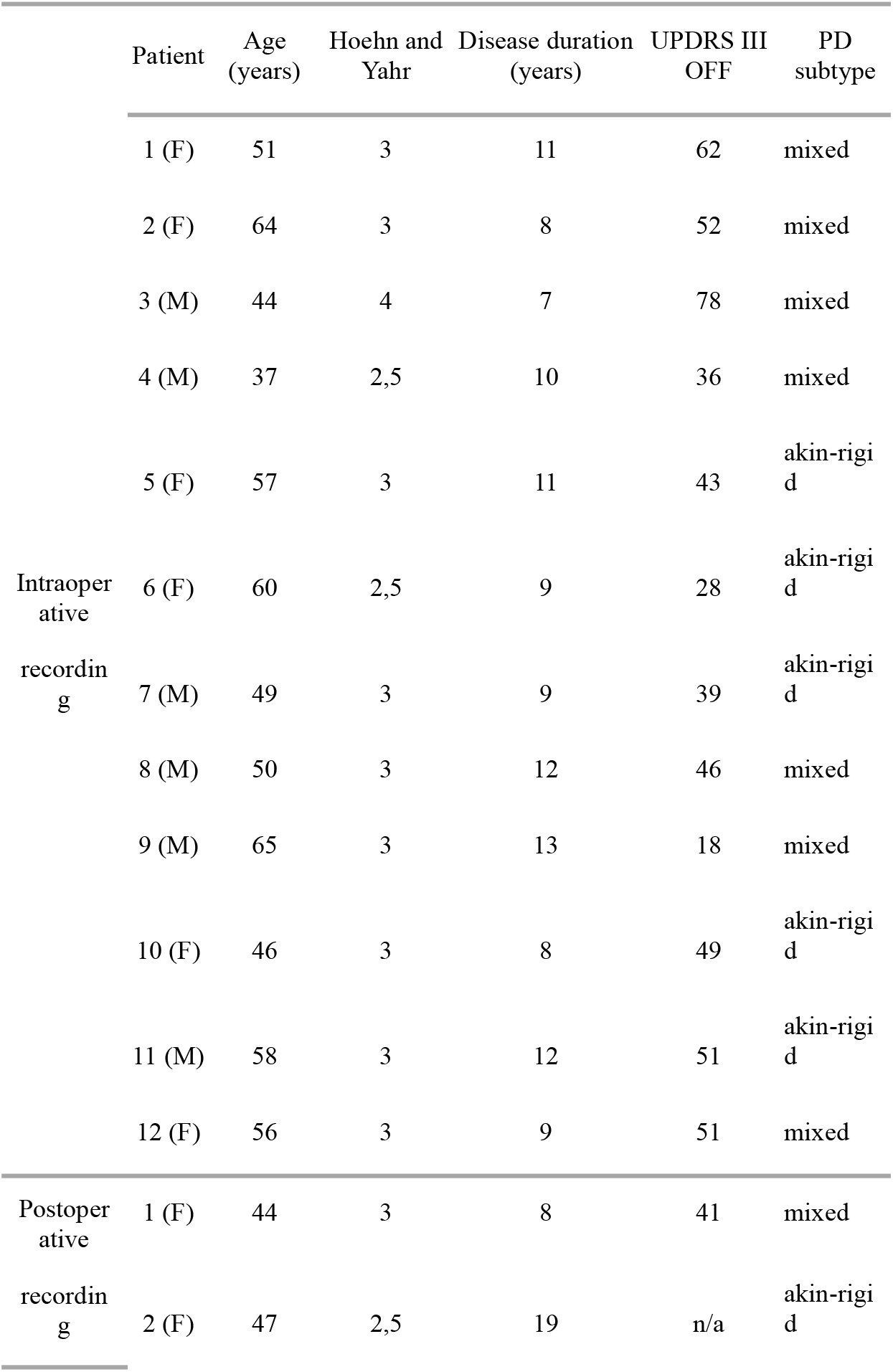
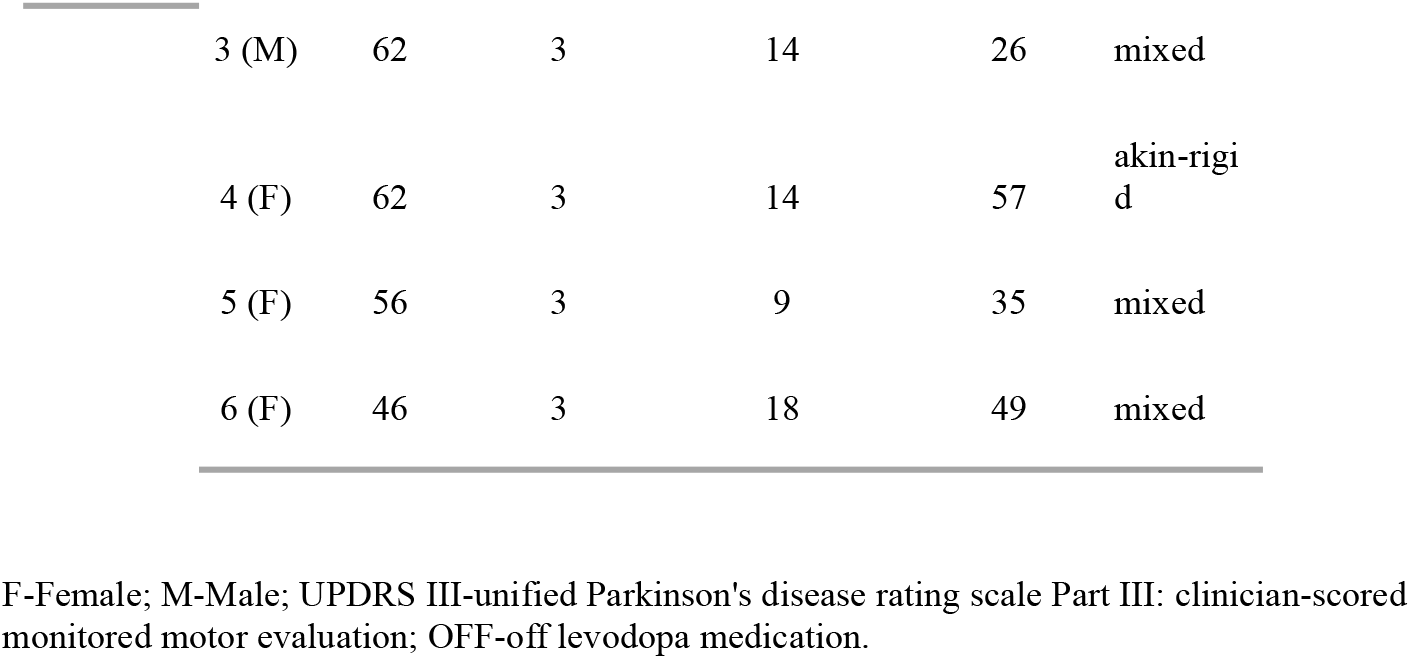
Clinical characteristics of Parkinson’s disease patients

The surgical procedure was consistent with the clinical standard-of-care. The data on the discharge activity of the STN neurons were obtained by microelectrode recording during planned stereotactic surgeries at the Burdenko Center of Neurosurgery. An informed consent was obtained from all subjects prior to their inclusion in the study. The study was approved by the Ethics Committee of the Burdenko National Medical Research Center of Neurosurgery. Microelectrode recording was used to identify the STN boundaries and select the optimal trajectory for deep brain stimulation (DBS) electrodes. The activity of neurons were recorded using a navigational NeuroNav system for inserting a microelectrode (Alpha Omega, Israel, www.alphaomega-eng.com), which was fixed to a stereotactic Leksell coordinate frame G (Elekta, Sweden) firmly affixed to the patient’s head. The calculated target point coordinates were: 3.5 to 4.5 mm below and 11.5 to 13 mm lateral to the CA-CP line, 1.5 to 2.5 mm posterior to the CA-CP midline. Neuronal activity was extracellularly recorded in the two brain hemispheres, starting at a distance of 15 mm to the calculated target point. The middle of the second contact of the DBS electrodes was placed in the middle of the STN, after refining its exact borders along the trajectory based on microelectrode recording (MER) data.

Along with MER, EMG signals for the flexor and extensor muscles of the hand were registered during the surgery with sampling rate 24 kHz. In the postoperative period (next day after surgery), a 16-channels recording of local field potentials from implanted DBS lead (St. Jude, USA) inside the STN along with EMG signals for the flexor and extensor muscles of the hand were recorded using the EEG system NeuronSpectrum65 (Neurosoft, Russia) with sampling rate 2 kHz. DBS leads were characterized by the length of one contact 1.5 mm, longitudinal spacing 0.5 mm and spatial configuration 1-3-3-1 (Model 6178). Only the upper and central bipolar signals were included in the study, since they were presumably located in the motor region of the STN (Jahanshahi et al., 2015).

During the registration of neural activity, patients were asked to perform movements driven by verbal commands of a researcher (external stimuli) as follows: “clench/unclench your right/left hand”. The duration of holding the fist was about 3-4 seconds, the number of repetitions varied from 5 to 10 times (6 ± 1.6) on each MER side. Also, patients were asked to clench/unclench their right / left fist without particular verbal commands at their own pace, the number of repetitions varied from 5 to 10 times (9 ± 3.3) on each MER side. During postoperative multichannel local field potentials (LFP) recordings, we asked patients to perform the same motor tests for 10 times.

### Data analysis

At the preprocessing stage, neurograms was filtered (100-3000 Hz) from interference and artifacts, and the activity of single neurons was sorted by the shape and amplitude of spikes, using the principal component analysis method in the Spike2 software (Cambridge Electronic Design, United Kingdom). EMG signals were bandpass-filtered (16-300 Hz) and rectified. The analysis of neural responses was performed by plotting the peristimulus rasters and the peristimulus spike histograms simultaneously with perievent histograms of rectified EMG signals using NeuroExplorer software (Nex, United States). Movement onset was determined manually using EMG traces. When choosing the epoch of analysis, we were guided by the duration of the IG movements (approx 0.5 sec), which was shorter than ET and also included the premovement period (0.5 sec). For each responding neuron, the amplitude of reactions, the duration of reactions and the latency period were evaluated.

The analysis of local field potentials included pre-filtering (0.5-300 Hz) of recorded signals and removing artifacts. To assess changes in beta activity, calculation of band (10-30 Hz) energy versus time was performed in NeuroExplorer software. The responses were analyzed by plotting peristimulus spectrograms, peristimulus histograms of the beta band energy and rectified EMG signals. The amplitude and duration of desynchronization of beta activity during ET and IG movements were evaluated.

Statistical analysis was performed using a nonparametric paired Wilcoxon test.

## 3 Results

Overall, we have found 26 neurons that were sensitive to assigned motor tests in twelve studied patients. We have categorized neuronal activity into 2 groups according to the types of responses: that is, activation (76.9%) and inhibition (23.1%). In 90% cases, an activation preceded the movement by 0.1-0.3 s, the magnitude of responses ranged from 16 to 109 spikes per second. Fifty-three and eight tenths percent of responses were tonic – they lasted throughout the entire motor act from clenching to unclenching a fist), while 46.2% appeared to be phasic – there were brief responses related to certain movement phase lasting for 0.1-0.5 s. In 15.4% cases, the STN neurons responded after the onset of the movement with the lag of 0.05 - 0.2 s and the response amplitude about 22-38 spikes per second. Sixty-six and seven tenths percent of inhibitory responses in the STN neurons were leading, occurring 0.2-0.3 s before the movement onset with the response amplitude about 10-31 spikes per second. Thirty-three and three tenths percent of inhibitory responses in the STN neurons were lagging, with time latency 0.1-0.2 s from movement onset and response amplitude ranging from 40 to 60 spikes per second. Eighty-three and three tenths percent of inhibitory neurons responded tonically and 16.7% responded in a phasic manner. All phasic responses preceded the movement. All responsive neurons that were studied here responded both to externally triggered and internally guided motor tests. In contrast to the more prominent and stable reactions during ET movements, the amplitude of responses to IG tests was reduced and in some cases attenuated gradually throughout the trial (Me Δfr ET = 33, Me Δfr IG = 26, p = 0.003 - Wilcoxon signed-rank test, see Figure 1).

**Figure 1.**
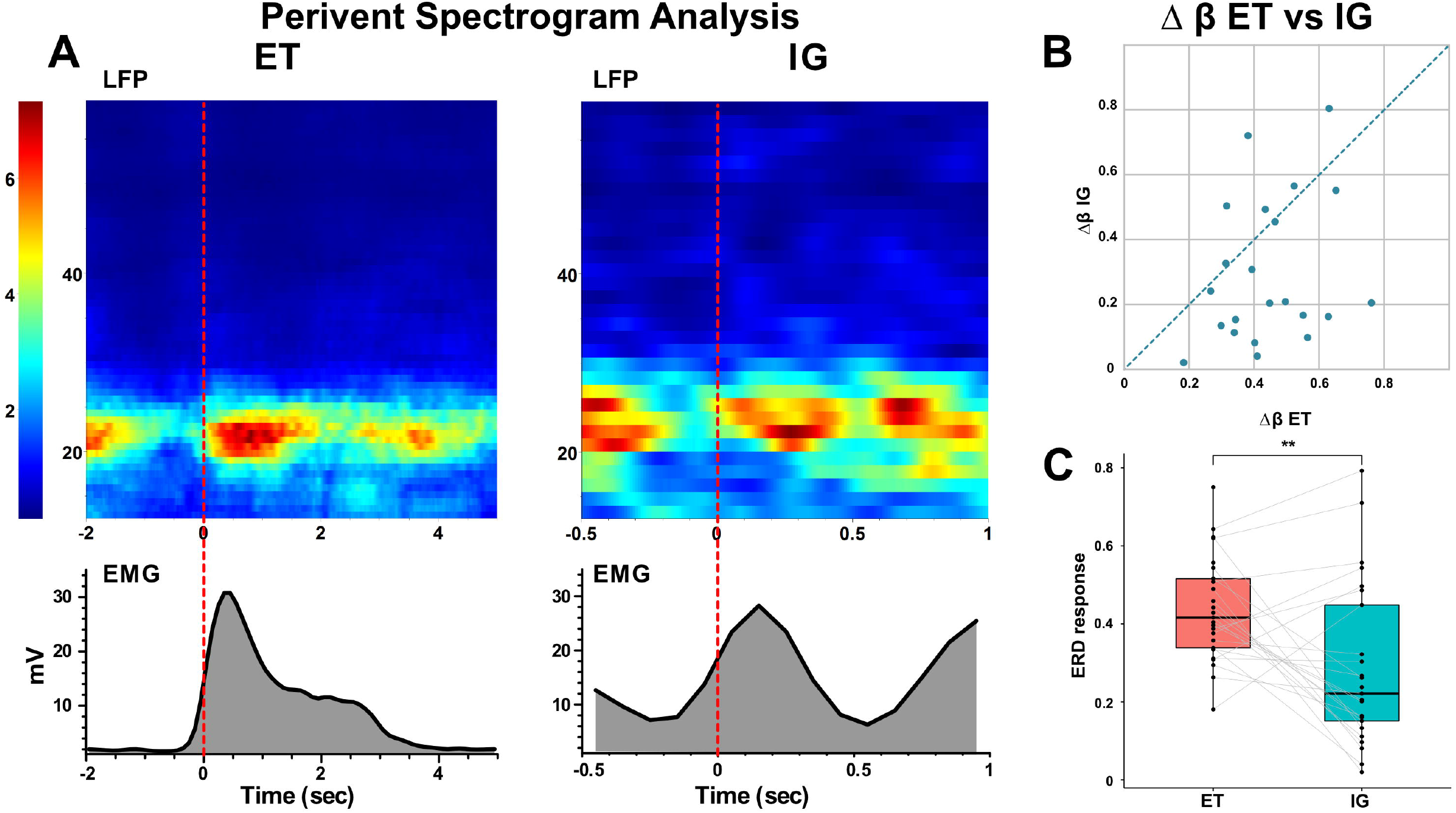
Neuronal responses of STN units to voluntary ET and IG movements in PD. A: an example of perievent raster plots, perievent spike histograms and perievent rectified EMGs for ET and IG movements. B: scatter plot of response amplitude (delta firing rate) to ET (x-axis) and IG (y-axis) movements. Each point marks data from one cell. C: boxplots with lines of cells responses to ET and IG movements.

Spectral analysis of populational neural activity (LFP) in the postoperative period showed a time-stable desynchronization beta activity during ET movements (i.e., the event-related desynchronization (ERD)) lasting 0.5-1 sec, which preceded the start of the movement by 0.1 - 0.4 seconds. There was also an event-related synchronization of the beta (i.e., the event-related synchronization (ERS)) after fist unclenching lasting for 1-1.2 seconds (Figure 2).

**Figure 2.**
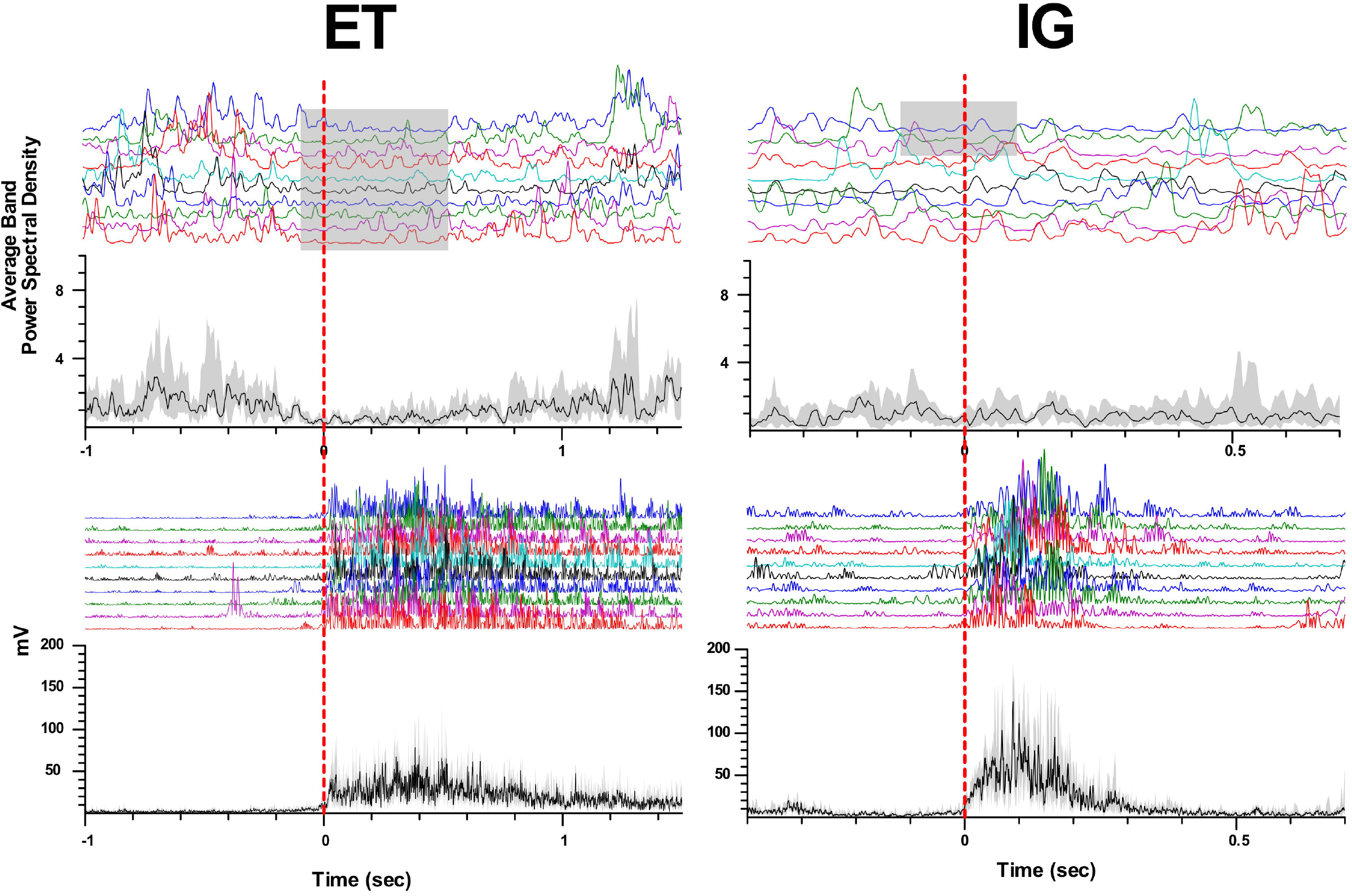
Event related beta desynchronization before and synchronization during voluntary ET and IG movements in PD. A: example of perievent spectrogram of LFPs and rectified EMGs during ET and IG movements. X-axis: time, sec; y-axis for LFP: frequency, Hz; y-axis for EMG: amplitude, mV; color scale shows spectrum power, %. B: scatter plot of beta ERD during ET (x-axis) and IG (y-axis) movements. Each point marks data from one bipolar. C: boxplots with lines of beta ERD to ET and IG movements.

When performing self-initiated movements, we observed the beta ERD preceding the initiation of the movement by 0.3-0.1 sec and lasting for 0.4-1 sec following beta ERS emerging after movement termination. This ERD response had decreased in amplitude in comparison with ET movements (Me Δβ ET = 0.42, Me Δβ IG = 0.22, p = 0.006 - Wilcoxon signed-rank test, see Figure 2). We found that in some cases ERD response to IG movement gradually faded after the first 2-3 trials of clenching the fist in contrast to stable ERD response to ET movement (Figure 3).

**Figure 3.**
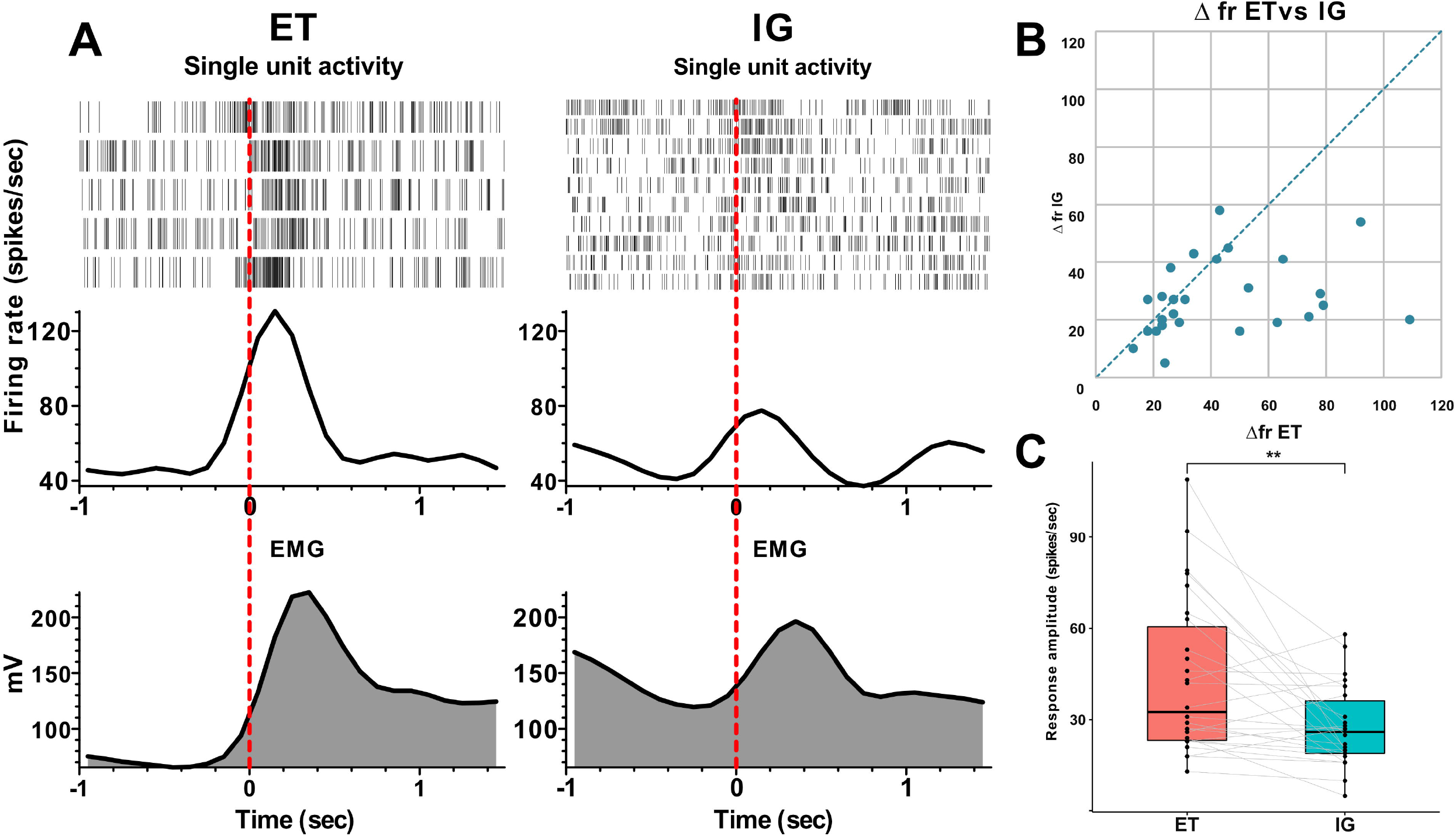
Attenuation of beta desynchronization during repeated IG movements in PD. A: examples of perievent raster and perievent histograms of beta energy power and rectified EMGs for ET and IG movements. X-axis: time, sec; y-axis: average beta energy power (a.u.) and rectified EMG amplitude (mV). Black line in perievent histograms shows median value, gray area shows interquartile range.

In some cases, we also observed tonic desynchronization of beta activity during IG movements, which started to recover gradually up to the level of background activity in the course of the movement. For some bipolars, prominent beta ERS was observed after the end of the block of motor tests for both ET and IG movements.

## 4 Discussion

### STN movement-related activity

According to the BG model, STN, being an integral part of the indirect and hyper-indirect pathway, inhibits unwanted movements (Alexander et al., 1986; Hasegawa et al., 2022). Recent studies of the single unit activity in monkeys have suggested that the STN may stabilize movements by reducing the variability of neural activity in the globus pallidus (Hasegawa et al., 2022). In a few studies of the STN single units in patients with PD, the authors have found neurons responding to movements of the hands, feet and face in the dorsolateral part of the nucleus (Rodriguez-Oroz, 2001; Abosch et al., 2002).

Here we described the dynamics of neural responses in the STN of patients performing motor tests. We revealed the polymodality of these reactions, which differed both in trend (activation or inhibition) and in response latency (lead or lag). These findings indicate a more complex functional role of the STN than the simple inhibition of movements predicted by the classical basal ganglia model (Alexander et al., 1986). The presence of advanced phasic neural responses suggests that the STNs may be involved in the formation of the motor programs and the initiation of voluntary movements or could be involved in leading cognitive aspects of motor control (Zavala et al., 2015; Aron et al., 2016). At the same time, tonic delayed responses indicate the contribution of the STN also to afferent motor control. These neural responses can be interpreted using a dynamic model of the basal ganglia, which predicts temporal and functional differences of the cortico-subthalamic and strio-pallido-subthalamic pathways (Nambu et al., 2002; Hasegawa et al., 2022). Moreover, recent animal studies have reported that initiation of movements may be performed primarily through parafascicular nucleus projections to STN, which were named by the authors as the superdirect pathway (Watson et al., 2021).

Analysis of the LFP in the STN also revealed different response dynamics during movement, i.e., advanced beta desynchronization followed by hypersynchronization throughout the motor execution. These results are in agreement with the previous data and indicate the important role of beta desynchronization for starting the motor program (Kühn et al., 2006; Little and Brown, 2014; Sharott et al., 2018). The variety of neural responses shown here and the dynamics of beta synchronization-desynchronization suppose an important functional role of the STN both in initiating the motor program through the cortico-subthalamic or thalamo-subthalamic pathways, as well as in contributing to the execution of the motor program through other intracerebral pathways (Polyakova et al., 2020).

### ET vs IG movement responses

According to the existing model, ET and IG movements are performed largely through segregated BGTM and CC paths (Cunnington et al., 2002; Cerasa et al., 2006; Taniwaki et al., 2006, 2013; Purzner et al., 2007; Hackney et al., 2015). Taking into account this model as well as the classical model of the basal ganglia (Alexander et al., 1986), we expected to detect neurons in the STN that would selectively respond to externally triggered movements but not internally guided ones. Contrary to our predictions, we failed to identify such neurons, instead we found that the responses of the STN neurons to self-initiated movements displayed a reduced amplitude and usually faded while motor tests were performed repeatedly.

Analysis of the populational neural activity inside the STN represented by LFP data revealed a stable ERD response with ET movements and a significant decrease in the amplitude of beta desynchronization with IG movements, as well as its gradual attenuation with repeated movements. The deficit of desynchronization in IG movement could be possibly explained by a smaller number of neurons involved in this type of movement, however, an analysis of the responses of single STN units disproved this assumption, as all examined units that responded to motor tests were sensitive to both types of movements.

An electrophysiological study by Bichsels et al., who also studied ET and IG movements from deep brain recordings in patients with PD, (Bichsel et al., 2018) described neural responses in the STN of PD patients that were similar to the ERD responses. The authors attribute these differences in responses to the existence of functionally segregated cortico-basal ganglia networks controlling motor behavior in PD patients and thus corroborate the previous assumption of PD patients being shifted from habitual towards goal-directed behavior (Bichsel et al., 2018). Authors showed tonic beta desynchronization during IG movement, but they didn’t study temporal dynamics of such responses. In some patients we found tonic beta desynchronization, which was attenuating during repeated movements. This observation should be proved in a larger sample of patients.

We assume that our results, on the one hand, indicate the participation of the basal ganglia in the selection of a relevant motor program and the initiation of both types of inspected movements. This may be possibly implemented through hyperdirect cortico-subthalamic projections or parafascicular nucleus (PF)-STN pathway associated with the regulation of attention and the selection of motor program (Nambu et al., 2002; Watson et al., 2021). On the other hand, these data emphasize the importance of an external stimulus that may assist the initiation of a motor program in patients with PD. Under the insufficient feedback in patients with PD, these external stimuli may contribute to the restart of the motor programs and, eventually, facilitate the execution of voluntary movements.

Significant beta desynchronization was found not only in the STN, but also in the globus pallidus and motor cortex of PD patients when performing ET and IG movements (Choi et al., 2020). This is consistent with the thesis that changes in beta activity in the BGTM network supporting movements are a prerequisite for normal motor control (van Wijk et al., 2017). However, the observed magnitude of beta ERD in the pre-movement period in GPi and PM/M1 was significantly greater for IG movements compared to ET movements. The differences in the desynchronization level in the GP found previously and in the STN presented here may be attributed to additional striopallidar projections entering the inner segment of the globus pallidus via a “direct” pathway.

In summary, we showed a decrease in the amplitude of neural responses of STN to IG movements performed repeatedly, as well as a reduced and gradually attenuating beta desynchronization when performing self-initiated movements in patients with PD compared with stable neural responses and beta ERD when performing ET movements. A probable explanation for this phenomena might be as follows. Presumably, each movement caused by an external signal may be viewed as a separate and complete motor program, which is restarted each time through verbal command and entailed beta desynchronization-synchronization. At the same time, in the case of self-initiated movements, the motor program is started once and under the impaired feedback afferentation observed in PD, the neural responses in the basal ganglia gradually fade. Our findings may contribute to a deeper understanding of the neural mechanisms causing the impairment of self-initiated movements in PD and indicate the value of an external stimulus for the implementation of voluntary motor behaviors in patients with PD.

## 5 Limitation

Our study has several limitations that should be mentioned. Intraoperative study of neuronal activity has a surgery time limitation and patients fatigue. This limits the number of motor tests presented to patients at each microelectrode recording site. А small number of patients does not allow correlation analysis between neuronal responses and clinical indicators of patients to prove that the observed results are associated with the disease. Due to ethical considerations, it is impossible to analyze neuronal responses in the basal ganglia in a group of healthy people.

This limitation will be partly compensated by studying the motor responses in the basal ganglia in patients with Parkinson’s disease in OFF- and ON-states and by comparison with data collected from cervical dystonia patients.

## Supporting information

Clinical characteristics of Parkinson`s disease patient

## Funding

The reported study was funded by RSF, project number 22-15-00344.

### Article types

Brief Research Report.

## Author Contributions

AS, VF contributed conception and design of the study. AT, SU, EB organized the database. VF, EB, AS performed data preprocessing, VF performed the statistical analysis. VF wrote the first draft of the manuscript. AS, EB edit manuscript. All authors contributed to manuscript revision, read, and approved the submitted version.

