## Supplementary material for "Attenuation of neural responses in subthalamic nucleus during internally guided voluntary movements in Parkinson’s disease": Clinical characteristics of Parkinson`s disease patient

Table 1. Clinical characteristics of Parkinson`s disease patients

|  | Patient | Age (years) | Hoehn and Yahr | Disease duration (years) | UPDRS III OFF | PD subtype |
| --- | --- | --- | --- | --- | --- | --- |
| Intraoperative  recording | 1 (F) | 51 | 3 | 11 | 62 | mixed |
|  | 2 (F) | 64 | 3 | 8 | 52 | mixed |
|  | 3 (M) | 44 | 4 | 7 | 78 | mixed |
|  | 4 (M) | 37 | 2,5 | 10 | 36 | mixed |
|  | 5 (F) | 57 | 3 | 11 | 43 | akin-rigid |
|  | 6 (F) | 60 | 2,5 | 9 | 28 | akin-rigid |
|  | 7 (M) | 49 | 3 | 9 | 39 | akin-rigid |
|  | 8 (M) | 50 | 3 | 12 | 46 | mixed |
|  | 9 (M) | 65 | 3 | 13 | 18 | mixed |
|  | 10 (F) | 46 | 3 | 8 | 49 | akin-rigid |
|  | 11 (M) | 58 | 3 | 12 | 51 | akin-rigid |
|  | 12 (F) | 56 | 3 | 9 | 51 | mixed |
| Postoperative  recording | 1 (F) | 44 | 3 | 8 | 41 | mixed |
|  | 2 (F) | 47 | 2,5 | 19 | n/a | akin-rigid |
|  | 3 (M) | 62 | 3 | 14 | 26 | mixed |
|  | 4 (F) | 62 | 3 | 14 | 57 | akin-rigid |
|  | 5 (F) | 56 | 3 | 9 | 35 | mixed |
|  | 6 (F) | 46 | 3 | 18 | 49 | mixed |

F-Female; M-Male; UPDRS III-unified Parkinson's disease rating scale Part III: clinician-scored monitored motor evaluation; OFF-off levodopa medication.
